# Cardiac Cell-Derived Matrices Impart Age-Specific Functional Properties to Human Cardiomyocytes

**DOI:** 10.1101/2020.07.31.231480

**Authors:** M. A. Kauss, S. J. Rockwood, A. C. Silva, D. A. Joy, N. Mendoza-Camacho, M. N. Whittaker, Erica Stevenson, N. J. Krogan, D. L. Swaney, T. C. McDevitt

## Abstract

Cell-derived matrices (CDMs) isolated from cultured cells provide complex and tissue-specific biochemical and physical cues derived from the extracellular matrix (ECM) that are lacking in typical tissue culture environments. However, current methods enhance ECM adhesion and thickness via introduction and promotion of singular matrix proteins, skewing the matrix composition, and confounding comparisons between CDMs. Here we developed a protocol that enhances CDM stability and deposition, respectively, by combining an L-polydopamine surface coating with Ficoll macromolecular crowing prior to hypotonic decellularization. This methodology was applied to the study of age-dependent phenotypic and functional changes observed in cardiac ECM by comparing the morphologic, electrophysiological and metabolic response of cardiomyocytes in response to CDMs produced by fetal and adult cardiac fibroblasts. Furthermore, mass spectrometry proteomics identified the enrichment of collagen VI in fetal CDMs, which we determined via siRNA-mediated silencing during CDM production to be necessary for maximal oxidative respiration in cardiomyocytes.

## Introduction

The dynamic and complex biochemical and physical cues provided by the extracellular matrix (ECM) significantly impact tissue function^1,2^, but have proven difficult to model *ex vivo. In vitro* cell biology studies often rely on ECM molecules as an adhesive for culture applications^3^, and de-emphasize their ability to serve as tissue-specific environmental cues. Many of the commonly used matrix coatings are comprised of individual, often recombinant, proteins^4^, whereas others, such as Matrigel, are derived from tumorigenic cells and offer limited tissue specificity^5^. Alternatively, *in vivo* studies of matrix biology that retain tissue complexity often contend with multiple confounding factors, such as preservation of organ structural integrity, prevalence of embryonic lethality, and heterotypic cell contributions to tissue microenvironment^6^.

Cell-derived matrices (CDMs) have emerged as a tractable experimental platform to model the ECM *in vitro*. CDMs are comprised of the isolated matrix produced by cultured cells – typically stromal cells from the tissue of interest^7,8^, which produce a large proportion of interstitial ECM^9^. After a suitable culture duration, stromal cultures can be decellularized while retaining the unique composition and structural organization of the extracellular matrix produced by the cells. The source, age, and genetic background of the cells can be manipulated to tailor CDM properties to model specific tissue systems and address biological questions. Additionally, CDMs are an exceptionally flexible platform to model ECM because they are adaptable to a variety of culture formats, are highly amenable to genetic perturbation, and limit the paracrine influence of viable cells that frequently confound alternative co-culture formats.

However, current methods for CDM matrix production and decellularization present caveats that limit CDM fidelity to mimic native ECM properties, with unknown influences on subsequent cell biology studies. Many protocols aim to improve the durability of ECM by promoting adhesion with pre-adsorbed and/or crosslinked matrix components, such as gelatin (denatured collagen) or fibronectin (FN)^7,10,11^, and increasing matrix assembly and thickness by promoting collagen I synthesis and secretion^8,10^. These types of approaches perturb the endogenous ECM composition by artificially increasing individual matrix proteins, sometimes to supraphysiological levels. Additionally, after ECM deposition, the method of decellularization can impact the composition and structure of the isolated matrix. Decellularization of tissues and CDMs often relies on the use of surfactants, such as SDS or Triton-X^7,10–12^, which strip away a large fraction of the growth factors, proteoglycans and glycosaminoglycans (GAGs) that are non-covalently bound to the matrix^13^, thereby significantly changing the relative composition of the ECM that had been produced. In addition, SDS has been shown to alter the stiffness and structure of collagens in decellularized tissues^14^. Thus, the susceptibility of CDMs to protocol-specific biases necessitates the need for careful methodological consideration in comparative matrix studies.

CDMs are particularly amenable to modeling and interrogating cell-ECM interactions at a structural and functional level, making them well suited to study cardiac biology. Cardiomyocyte (CM) behavior relies heavily on morphology and signaling events^15–20^, which are both influenced and reinforced by the local ECM microenvironment^6,21,22^. Specifically, the fetal cardiac ECM is loosely organized^15,23^ and enriched with growth factors and GAGs^24,25^ to support the developing myocardium and smaller, proliferative fetal CMs^19,26–28^, whereas the adult ECM is a more fibrillar^23^ and collagen-rich^29^ environment intended to main homeostasis of highly aligned, energetically-demanding adult CMs that constantly experience significant contractile strain^30–32^. The structural and compositional complexity of these age-specific cardiac ECMs are challenging to model *in vitro*. Historically, the majority of cardiac ECM studies have either been performed *in vivo*, which is complicated by embryonic lethality^6^, or with tissue ECM isolated using methods that either partially or completely denature the matrix structure and remove non-covalently bound components^29^. Thus, a robust protocol for CDM production is crucial to enable more mechanistic *in vitro* studies of the structural and compositional complexity of cardiac ECM in simulated native physiological conditions.

This study aimed to determine how cardiac ECM at different developmental stages impacts the physiology and function of CMs. To accomplish this, we developed a novel method for cardiac CDM production from fetal and adult primary cardiac fibroblasts (CFs) that avoided use of exogenous matrix proteins and detergents. Transcriptomic and proteomic analysis, before and after decellularization, respectively, revealed age-specific differences in CDMs that affected the function of CMs derived from induced pluripotent stem cells (CMs). In particular, CMs responded to the organizational and compositional differences in age-specific CDMs by adapting distinct morphological, electrophysiological and metabolic profiles, which we interrogated by genetic perturbation of CDM composition. Thus, our CDM model enables a detailed investigation of age-specific cardiac ECM properties, and the direct correlation of CM morphological and functional responses to better define ECM influences on dynamic functional shifts exhibited by cardiomyocytes during distinct stages of development.

## Results

### Development of Cell-Derived Matrices

CFs synthesize and organize the majority of the ECM in the heart^9^. Therefore, to model the cardiac ECM, we produced CDMs using primary cardiac CFs. The addition of exogenous matrix proteins and ascorbic acid (normally supplemented to promote collagens) were omitted during CDM production, and cultures were decellularized by established detergent-free methods^33^ (Fig. 1A) to preserve the native ECM composition. In the absence of commonly-used substrate coatings, such as crosslinked fibronectin or gelatin, tissue culture plates were pre-treated with L-polydopamine (dopa)^34,35^, an inert biopolymer that promotes matrix retention in a concentrationdependent manner through covalent binding of amine groups that are abundant in matrix proteins. Dopa concentrations ranging from 0.025 – 2 mg/ml were pre-adsorbed onto tissue culture plastic (TCP) prior to murine CF seeding, and cultures were then decellularized via hypotonic bursting after 10 days (Fig. 1B). Matrices formed equally on all dopa concentrations, but at 0.2mg/ml dopa and lower, decellularization induced partial or complete loss of CDM, resulting in the folding or absence of the matrix layer. The highest concentration tested, 2mg/ml, reliably retained mouse (Fig. 1B) and human (Fig. 1C) CDMs throughout the decellularization process, and thus was used for all subsequent studies. Secondly, to enhance matrix deposition and the resulting CDM thickness, we added macromolecular crowders (MMCs) to fibroblast culture, which reduce the effective volume for molecule diffusion, and thereby increase the effective concentration of soluble molecules^36^. MMCs increase cell-matrix associations and reduce diffusion-associated loss, which promotes remodeling of secreted proteins and other molecules into the local ECM. MMC culture with the neutrally charged Ficoll (8.5:1 molar ratio of 70kDa and 400kDa), promoted alignment of human CFs (Fig. 1C), which resulted in greater alignment of the matrix fibrils in the CDMs, as demonstrated by colorimetric labeling of FN fibril orientation post-decellularization (Fig. 1D). Additionally, supplementation of the media with Ficoll increased ECM abundance, as determined by CDM dry weight (Fig. 1E) and thickness (Fig. 1F). While the confocal microscopy necessary to perform high quality optical sectioning on CDMs required CDMs to be produced on glass, (Fig. 1F), CDMs produced on TCP appeared notably thicker during optical sectioning, both with and without MMCs (Supp. Fig. 1A, B). Other known MMCs, dextran sulfate and polystyrene sulfonate^37^, induced cytotoxicity at the doses tested (Supp. Fig. 1C), and therefore were not pursued further. The combination of dopa pre-coating and CF culture with Ficoll as an MMC enabled the production of robust CDMs that withstood decellularization without the use of exogenous matrix factors, as demonstrated with human CDMs in Fig. 1G.

**Figure 1.**
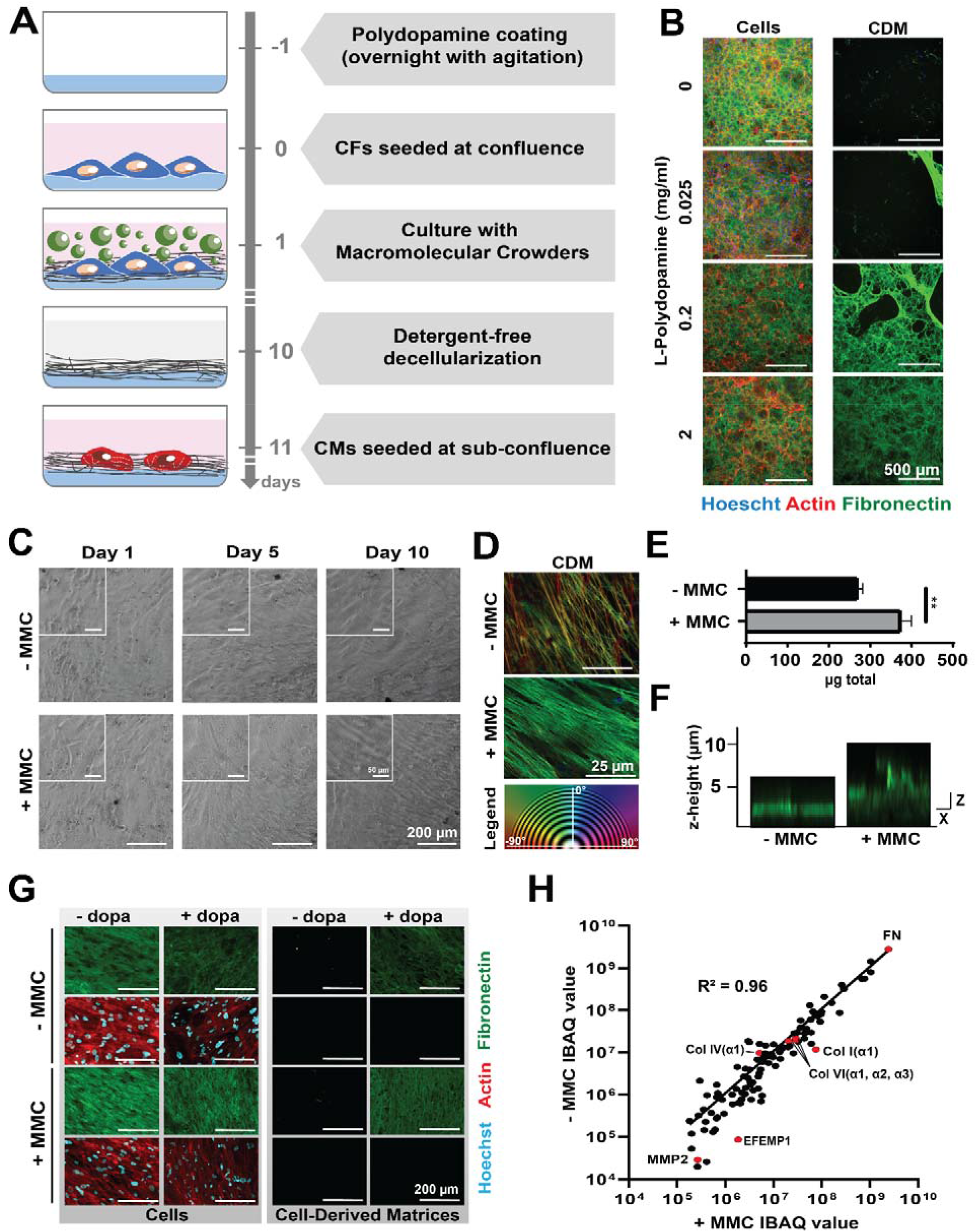
Production and characterization of cell-derived matrices (CDMs). A) Overview of CDM production and use. B) Decellularization of murine CDMs on escalating concentrations of L-polydopamine demonstrate concentration-dependent retention. C) Brightfield imaging of human CFs exhibit increased alignment over culture duration with Macromolecular Crowders (MMCs). D) Alignment of fibronectin fibrils in human CDMs produced with and without MMCs, pseudocolored by fibril orientation. E) Dry weight of human CDMs. ** p < 0.01. F) Human CDM thickness demonstrated by optical sectioning via confocal microscopy. G) Fluorescent microscopy of human CFs before and after decellularization protocol. H) Correlation of individual proteins intensity-based absolute quantification (IBAQ) within human CDMs produced with and without MMCs, quantified by liquid chromatography tandem mass spectrometry (LC-MS/MS).

To assess the influence of MMC addition on CDM composition, the matrix protein content of human adult CF-derived CDMs produced with and without MMCs was assayed by liquid chromatography with tandem mass spectrometry (LC-MS/MS)^38^. Linear regression analysis of protein abundance revealed a Pearson’s correlation coefficient of R^2^ = 0.96 between the two conditions (Fig. 1H), indicating that MMCs did not drastically alter the native matrix composition. FN, the foundational matrix protein required for binding and organization of many other matrix proteins^39,40^, was the most abundant protein identified in CDMs, and present at the same proportion regardless of the presence of MMCs, as was the basement membrane protein Collagen IV, alpha chain 1. Very few proteins did not exhibit a linear correlation between conditions; some, such as Col I and matrix metalloproteinase 2 (MMP2), were increased three-to-five fold with the addition of crowders. The increase of MMP2 in response to MMCs reported previously by transcriptional analysis^41^ may reflect more active remodeling required to produce thicker CDMs. These data demonstrate that while MMCs increase the thickness and abundance of ECM and matrix-associated proteins, the majority of the ECM composition remains unaltered. Thus, the combination of dopa pre-coating and MMCs enabled the production of CDMs without exogenous matrix factors, thereby providing a platform for unbiased comparative studies of matrix biology.

### Age-Specific Cardiac CDMs

*Ex vivo* modeling provides an opportunity to dissect how specific components of the ECM influence cardiac function, particularly as the heart undergoes significant developmental and physiological changes^2,6,15^. To create human cardiac matrices reflective of differing developmental stages, we produced CDMs from both fetal and adult human primary CFs. CFs of both ages adopted cellular alignment over 10 days of culture with MMCs (Fig. 2A), resulting in aligned CDMs (Fig. 2B,C). Fetal CFs, which are smaller and more proliferative than adult CFs, quickly achieved alignment by day 5 of culture with MMCs, whereas adult CFs were not fully aligned until closer to day 10 (Fig. 2C). The resulting fetal and adult CDMs also exhibited distinct ultrastructure. Specifically, fetal CDMs were more loosely organized with coiled fibrils, whereas adult CDMs exhibited more highly aligned and bundled matrix fibrils (Fig. 2D), which correspond to age-dependent structural differences observed in decellularized murine ventricular tissue^23^. Furthermore, CDM GAG content was preserved through decellularization, and was more enriched in fetal CDMs than adult (Fig. 2E), as expected^42^. GAG abundance was not homogenous within CDMs, but instead displayed regional differences dependent on CF growth patterns, resulting in a swirled pattern in fetal CDMs and more diffuse presence throughout adult CDMs. However, both fetal and adult matrices retained demonstrably more GAGs than Matrigel (Suppl Fig. 2), a common culture substrate for CMs derived from induced pluripotent stem cells (iPSC-CMs).

**Figure 2.**
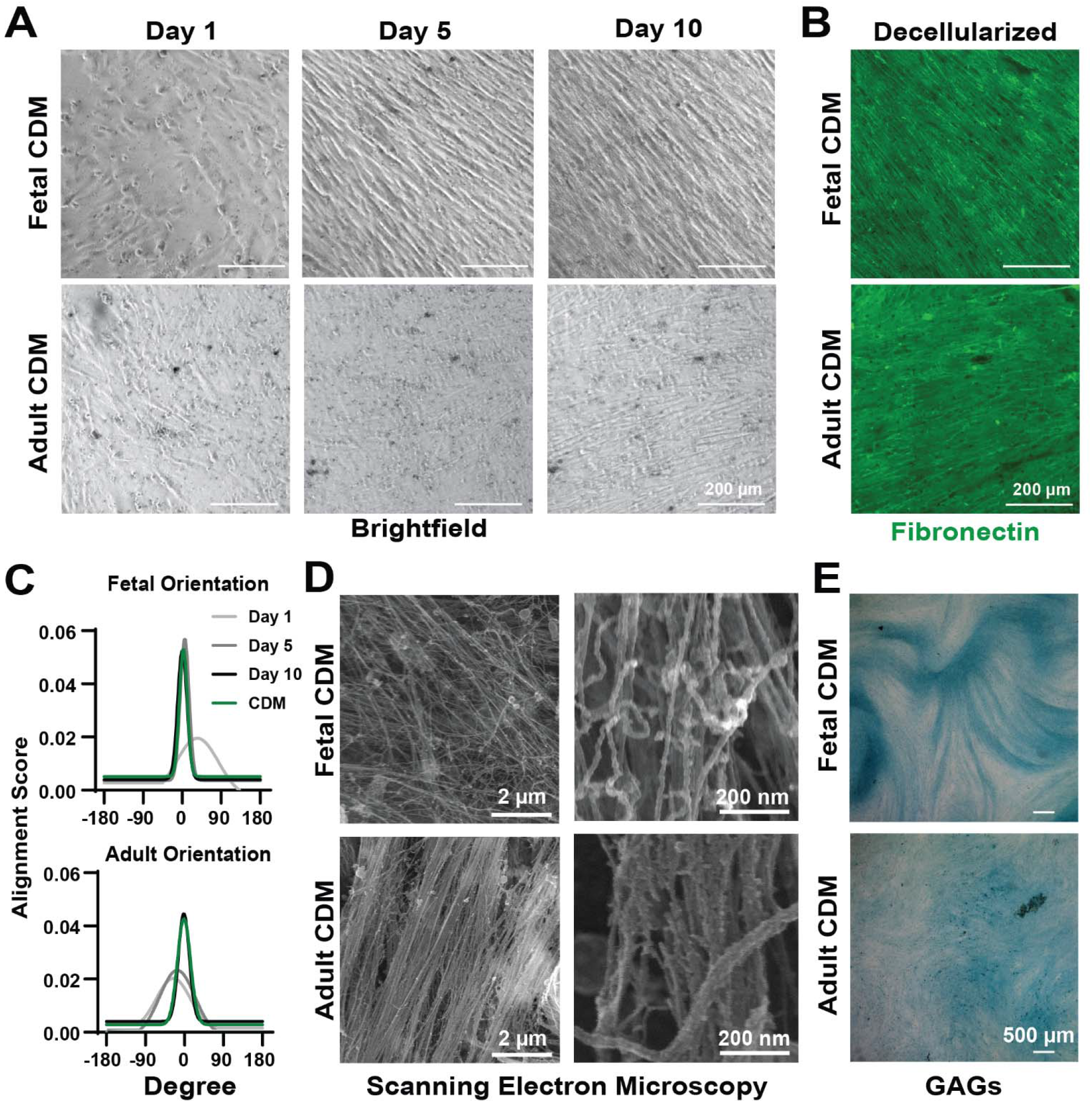
Fetal and adult human CDMs. A) Longitudinal brightfield imaging of fetal and adult CFs during CDM production. B) Resulting CDMs visualized by immunofluorescent labeling of fibronectin. C) Alignment of CFs and resulting ECM during CDM production. D) Ultrastructure of CDM fibrils produced by age-specific CFs detected by scanning electron microscopy. E) Glycosaminoglycans (GAGs) retained in CDMs post-decellularization, detected by alcian blue staining.

Age-specific compositional differences were assessed by longitudinal transcriptional analysis throughout CF culture and by proteomic analysis following decellularization of CDMs. In CFs, the geometric mean of transcript expression assayed at days 0, 5 and 10 of culture revealed distinct temporal patterns of key genes among the 496 matrix and matrix-associated genes (Supp Fig. 3). Genes encoding crucial matrix proteins responsible for the structural integrity and adhesion of the ECM (foundational genes) were expressed consistently throughout culture in both fetal and adult CFs. However, expression of proteases and pro-mitotic growth factors, such as neuregulin (NRG1)^43^, were highly enhanced in fetal CFs over time in culture, consistent with the growth factor-rich environment of the fetal, regenerative heart^24,43^, whereas adult CFs displayed increased expression of inflammatory cytokines, including IL-1β, which is expected of adult matrix-producing CFs^44^. Of the 419 matrix genes commonly expressed by fetal and adult CFs, only 98 (23%) were detected at the protein level in both fetal and adult CDMs (Fig. 3A). The incomplete overlap of transcriptional and proteomic results is likely due to the transient nature of gene expression and inevitable loss of some matricellular and non-covalently bound proteins during decellularization. In total, 133 proteins were detected in decellularized CDMs by LC-MS/MS, with 109 proteins common to CDMs of both ages. The common proteins represented all major categories of fundamental matrix and matrix-associated proteins (Fig. 3B), as annotated by the Matrisome Project^45^. CDMs from both fetal and adult CFs contained foundational matrix proteins, including FN, Col I, Col IV, and laminins, with no significant agespecific differences in abundance. There were, however, compositional features unique to agespecific CDMs. Nineteen proteins were significantly increased greater than two-fold in fetal CDMs relative to adult CDMs, and 15 were significantly decreased (Fig. 3C). GO enrichment analysis via the STRING^46^ database identified fetal CDMs as enriched for collagen fibril organization and growth factor binding, whereas adult CDMs were enriched for peptide crosslinking and calcium-dependent protein binding (Table 1); differences which reflect key ontogenic characteristics consistent with mammalian cardiac matrix development, and the structural organization of matrix fibrils observed in fetal and adult CDMs (Fig. 2D). Similar to fetal cardiac ECM^27,47–49^, fetal CDMs were enriched for fibrillin 2 and tenascin, which can influence morphogenesis, and developmental factors involved in cardiac specification, such as Wnt5a^21^. Adult CDMs, on the other hand, were enriched for several matrix-modulating proteins that promote matrix abundance, crosslinking and stabilization, such as transglutaminase 2^50,51^, hyaluronan and proteoglycan link protein 1^6,52^, transforming growth factor β2^53^, latent TGFβ-binding protein 2^54^ and insulin-like growth factor binding protein 7^55^ (Fig. 3C). The enrichment of these factors in adult CDMs is consistent with adult cardiac biology, which exhibits increased ECM crosslinking^56^ and organization^23^ and is associated with greater CM maturity^22,57^.

**Figure 3.**
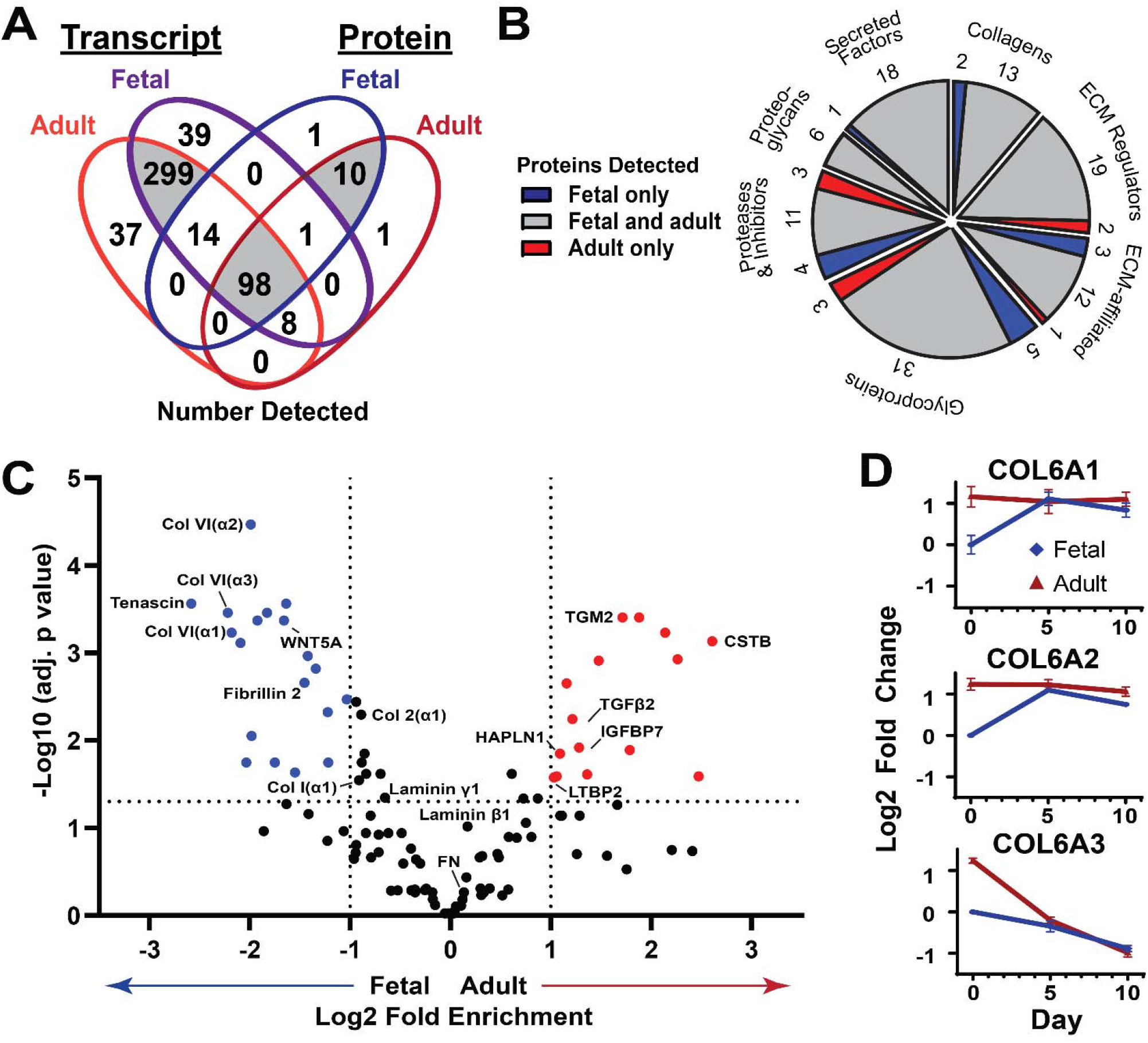
Age-specific transcripts and proteins for fetal and adult human cardiac matrices. A) Comparative Venn diagram of matrix-associated transcripts and proteins in fetal and adult CFs and CDMs, detected by RNA-Seq and LC-MS/MS. B) Proteins detected in fetal CDMs (blue), adult CDMs (red) or both (grey), per matrix protein categorization. C) Volcano plot of enrichment of individual matrix proteins. D) Temporal transcript expression of Collagen 6 chains in agespecific CFs.

**Table 1.** GO enrichment analysis of fetal and adult human CDM protein composition by STRING.

| CDM | Biological Process |  |  | Molecular Function |  |  |
| --- | --- | --- | --- | --- | --- | --- |
|  | *GO term* | *Description* | *FDR* | *GO term* | *Description* | *FDR* |
| Fetal | GO:0030198 | extracellular matrix organization | 5.93E-29 | GO:0019838 | growth factor binding | 8.63E-08 |
|  | GO:0009653 | anatomical structure morphogenesis | 5.85E-14 | GO:0005201 | extracellular matrix structural constituent | 1.80E-06 |
|  | GO:0048646 | anatomical structure formation involved in morphogenesis | 2.41E-11 | GO:0048407 | platelet-derived growth factor binding | 1.80E-06 |
|  | GO:0030199 | collagen fibril organization | 2.90E-11 | GO:0005102 | signaling receptor binding | 1.63E-05 |
|  | GO:0072359 | circulatory system development | 2.25E-09 | GO:0005178 | integrin binding | 1.63E-05 |
| Adult | GO:0032940 | secretion by cell | 0.00044 | GO:0048306 | calcium-dependent protein binding | 3.37E-14 |
|  | GO:0018149 | peptide cross-linking | 0.00073 | GO:0005509 | calcium ion binding | 6.48E-12 |
|  | GO:0045055 | regulated exocytosis | 0.001 | GO:0005544 | calcium-dependent phospholipid binding | 4.31E-09 |
|  | GO:0009888 | tissue development | 0.0013 | GO:0044548 | S100 protein binding | 3.37E-07 |
|  | GO:0016192 | vesicle-mediated transport | 0.0016 | GO:0046872 | metal ion binding | 1.19E-05 |

Since all three primary Col VI chains (Col 6) were enriched in fetal CDMs, we assessed the temporal transcriptional data to determine whether protein enrichment was due solely to increased expression of Col 6 genes in fetal CF. However, although fetal expression of Col 6 chains increased over the time in culture, transcript abundance was less than or equal to that of adult expression at days 0, 5, and 10 of CF culture (Fig. 3D). Due to the lack of correlation between transcript and protein abundance for Col 6, we investigated whether the retention of Col 6 in fetal CDMs was due to other factors enriched in fetal CDMs, potentially including its binding partners, Col I, Col IV, laminin^58^ and FN^59^. Of these foundational matrix proteins, only Col I appeared to be significantly enriched, and only mildly (Fig. 3C). It is therefore possible that subtle differences in Col I abundance contributed to enhanced retention of Col 6 in fetal CDMs. However, this does not rule out the potential that Col 6 retention is influenced by other factors such as specific temporal matrix dynamics, matrix organization, or non-protein components of the CDMs.

### iPSC-CM Response to Age-Specific Cardiac CDMs

To determine the influence of age-specific differences in CDM content and structure on human CMs, we cultured iPSC-CMs on fetal and adult cardiac CDMs. Confocal imaging revealed that CMs embedded themselves into the CDMs, enabling 3D matrix interactions instead of the basally-restricted matrix cues provided by Matrigel (Fig. 4A, XZ and YZ views). Relative to CMs cultured on Matrigel, CMs grown on CDMs exhibited increased cellular alignment, as visualized by FN and cTnT (Fig. 4B) and desmosome-associated protein plakophilin-2 via immunocytochemistry (Fig. 4C). The anisotropic enrichment of plakophilin-2 at the putative intercalated disks between CMs, which promotes electrophysiological coupling in maturing CMs^60^, was most apparent in CMs cultured on adult CDMs (Fig. 4C).

**Figure 4.**
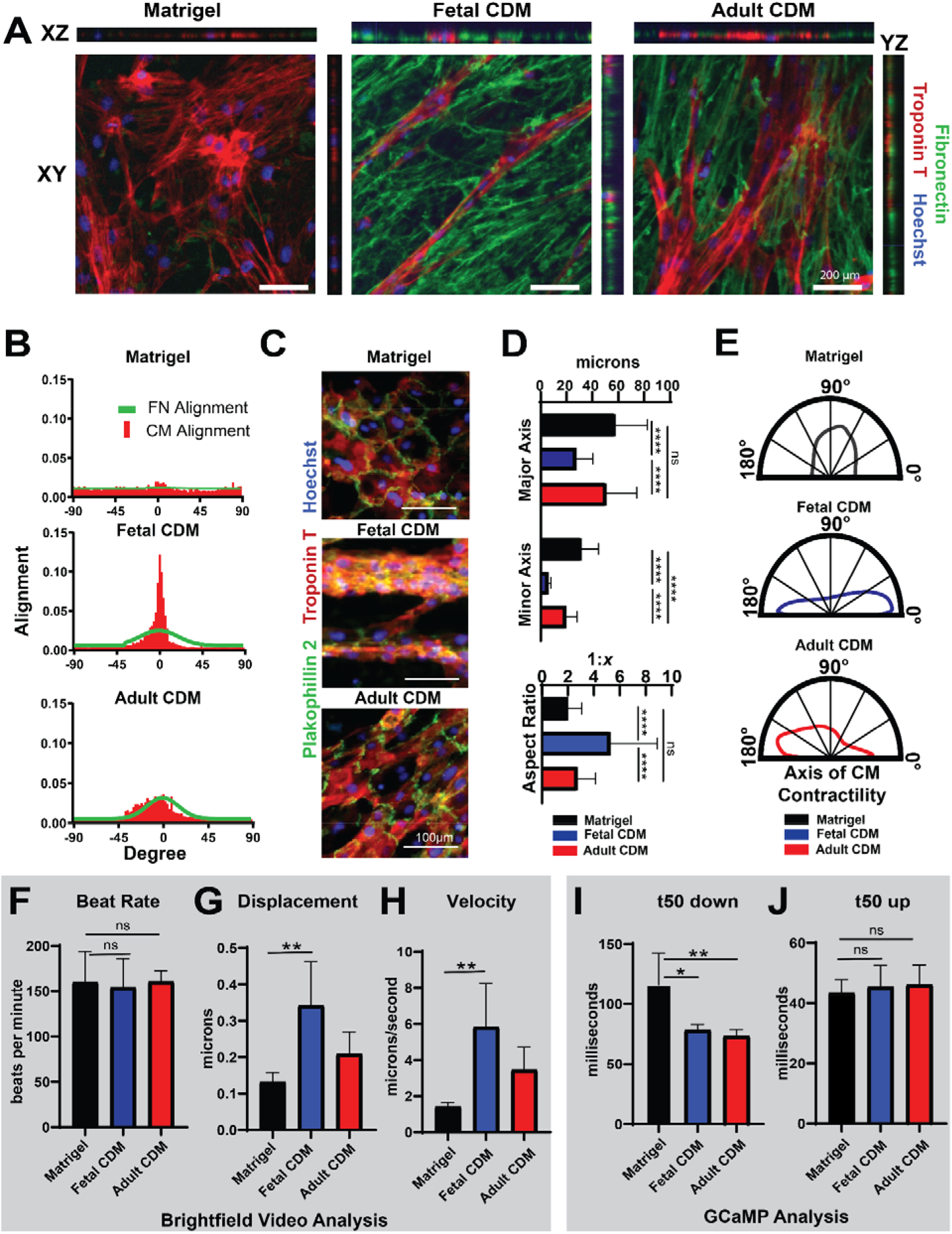
Human iPSC-cardiomyocytes (CMs) cultured on Matrigel and fetal and adult CDMs. A) iPSC-CM morphology when cultured in XY, XZ and YZ planes. B) Alignment of fibronectin fibers in Matrigel and CDMs, and alignment of corresponding cultured iPSC-CMs. C-D) Individual cell morphology and adherens junctions of iPSC-CMs, labeled by plakophilin 2 (C), and quantification of cell size and aspect ratio (D). E) Angle of iPSC-CM contractility, determined by live video analysis. F-H) iPSC-CM beat rate (F), displacement (G) and velocity (H) determined by live brightfield video analysis. I-J) Calcium peak time to decay by 50% (I) and time to rise by 50% (J) determined by live video analysis of genetically encoded GCaMP calcium reporter. ns = not significant, * p < 0.05, ** p < 0.01, **** p < 0.001.

CM function is also strongly influenced by morphology^22^, with cells increasing in size as they mature, and high aspect-ratio CMs achieving the most efficient contractions^16^. CMs cultured on fetal CDMs exhibited greater than a two-fold increase in morphological aspect ratio than those cultured on Matrigel, and CMs on both fetal and adult CDMs adopted narrower cell widths than those on Matrigel (Fig. 4D). Aligned culture on CDMs also had a marked effect on CM contractility and calcium handling. CMs grown on fetal and adult CDMs exhibited much more uniform axes of contraction compared to those cultured on Matrigel, as determined by optical flow analyses (Fig. 4E). Live-imaging and analysis of spontaneous beating showed that the CDMs did not impact CM beat rate (Fig. 4F), but culture on fetal CDMs increased CM contraction velocity (Fig. 4G), which resulted in three-fold greater total displacement (Fig. 4H). Additionally, relative to CMs cultured on Matrigel, calcium handling analyses indicated that iPSC-CMs grown on both fetal and adult CDMs had a much faster downstroke (t50 down), which is regulated by potassium and calcium channels (Fig. 4I), with no change in the upstroke (t50 up) that is regulated by sodium channels (Fig. 4J)^61^.

Collectively, these data indicate that CDMs improve the structural and contractile function of CMs by promoting alignment, contraction velocity, and specific calcium handling properties that more closely resemble native heart tissue than those achieved by CMs cultured on Matrigel.

### Collagen 6 Impacts on CM Respiration

One of the most marked age-specific compositional differences between CDMs was the fetal enrichment for Col 6. In skeletal muscle, Col 6 is crucial to metabolic function, and its loss induces metabolic dysregulation that promotes muscular dystrophies and reduces regeneration^62,63^. Though cardiac dysfunction in these conditions is rare, efforts to investigate the similarities between skeletal muscle and cardiac response to Col 6 deficiencies *in vivo* have been masked by non-CM phenotypes^64^. Therefore, we used CDMs as a platform to investigate the CM metabolic response to the loss of individual matrix proteins in complex, structurally-relevant ECM. To produce CDMs lacking Col 6, adult and fetal CFs were transfected with nontargeting (NT) siRNAs, or a combination of siRNAs inhibiting COL6A1, COL6A2 and COL6A3 (KD) during CF culture. Since CDMs represent the accumulation of protein production, Col 6 was assessed in CFs after 10 days of culture with siRNA silencing by western blot (Fig. 5A), and in decellularized CDMs by immunofluorescent labeling (Fig. 5B). These analyses confirmed Col 6 enrichment in non-silenced fetal CDMs, and showed that Col 6 protein was dramatically reduced following Col 6 KD. In several cases, suppression of Col 6 destabilized the ECM to the point of CDM loss, and this was particularly frequent in the fetal CDMs (Supplemental Fig. 4A). The fetal CDMs that remained intact contained some residual Col 6 protein, suggesting that the adult CDM, which innately has less Col 6, is better able to compensate for absence of this protein. In CMs cultured on fetal and adult CDMs, Col 6 KD had no impact on CM contractile properties, including beat rate, contractile displacement velocity, or downstroke time (Fig. 5C-F), which is consistent with the lack of reported cardiac dysfunction in Col 6-dependent muscular dystrophies^65^.

**Figure 5.**
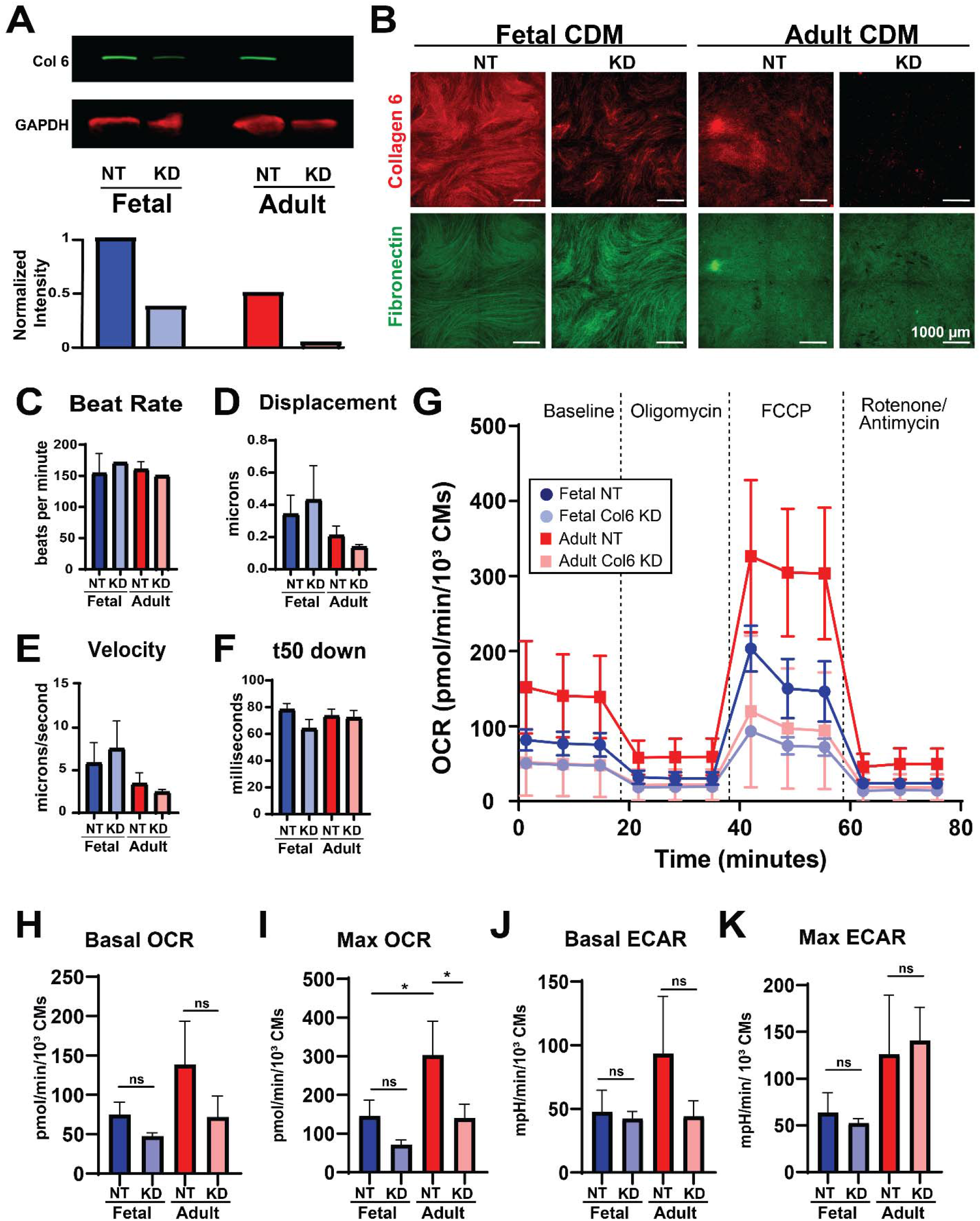
Collagen 6 knock down in CDMs. A) Western blot and normalized quantification of collagen 6 (Col 6) in fetal and adult CDMs after transfection with non-targeting (NT) or anti-Collagen 6 (KD) siRNAs. B) Immunocytochemistry visualization of Col 6 after KD in CDMs. C-F) Contraction characterization of iPSC-CMs cultured on CDMs with Col 6 KD, determined by live brightfield video analysis (C-E) and live GCaMP video analysis (F). G-K) Oxygen consumption rate (OCR) and extracellular acidification rate (ECAR) of iPSC-CMs cultured on NT and KD fetal and adult CDMs. G) Representative OCR measurements on Seahorse XF. H-K) Quantification of basal (baseline) and maximal (FCCP-treated) OCR and ECAR. ns = not significant, * p<0.05.

To characterize the influence of Col 6 on CM metabolism, CDM production was adapted to a 96-well plate format compatible with the Seahorse Extracellular Flux Analyzer. CM oxygen consumption rate (OCR) was assessed as a surrogate measurement for oxidative phosphorylation^66^, an indicator of CM metabolic maturity^67^, and extracellular acidification rate (ECAR) was assessed as measure of glycolysis^66^. After 3 days of culture, CMs grown on adult CDMs had a higher maximal OCR than those on fetal CDMs (Fig. 5G, I), which is consistent with the increased oxidative phosphorylation observed in maturing CMs in the adult heart. Additionally, Matrigel only supported OCR and ECAR levels similar to that of fetal CDMs (Supp Fig. 4). Col 6 KD in fetal CDMs did not significantly alter OCR or ECAR properties, potentially due to the presence of residual Col 6 protein. However, Col 6 KD in adult CDMs reduced the maximal OCR achieved by CMs (Fig. 5G, I), without significantly altering basal OCR or ECAR (Fig. 5G, H, J, K). The loss of Col 6 from adult CDMs, thus, influences CM metabolic respiration, specifically during simulated high energy demand, which is essential for metabolically active CMs to mature and respond to functional stressors and challenges. Therefore, genetic suppression of Col 6 had distinct influences on age-specific CDMs, which directly influenced CM metabolism.

## Discussion

The use of CDMs to study fundamental principles of cell-matrix interactions is an attractive approach that has increasingly grown in popularity in recent years^8,10,41,68^. CDMs are readily amenable to genetic and chemical modification, and provide a newfound opportunity to study complex ECM interactions without the inherent risks of *in vivo* knockout studies, such as embryonic lethality and confounding paracrine signaling. To expand the utility of CDMs, we sought to model age-specific characteristics of heart development reflected by the cardiac ECM using fetal and adult cardiac fibroblasts. Pre-treatment of tissue culture surfaces with the bioadhesive L-polydopamine covalently stabilized extracellular matrices following decellularization, while circumventing the need for adsorption of exogenous matrix proteins or complex surface chemistries and/or crosslinking. In addition, although MMCs have been used previously to enhance overall matrix production, characterization of MMC effects has been primarily limited to a few canonical ECM proteins such as Col I, FN and MMP2^41^. We show that the matrix produced in the presence of the MMC Ficoll is highly correlated with composition of matrix derived by CFs in the absence of MMCs. The combination of the dopa/MMC strategy in CDM models enabled the isolation of exclusively cell-produced matrix, which is crucial to studies involving inherent ECM comparisons between distinct physiological conditions. Under these conditions, fetal and adult CDMs retained a number of critical age-specific features of ECM structure and composition which differentially impacted iPSC-CM morphology and function. Although fetal and adult CDMs both promoted CM alignment, fetal CDMs promoted greater CM contraction velocities whereas adult CDMs promoted anisotropic desmosome enrichment at CM junctions - a feature that is associated with increased maturity. These differential responses to age-specific CDMs reflect the inherent influence of the ECM on cardiac physiology, as these differences were observed after a relatively short culture duration, in contrast to the extended culture that is often required to induce demonstrable maturation of stem cell-derived CMs on minimal ECM substrates^69^. Importantly, CDMs – especially the fetal CDM – retained a notable fraction of GAGs, which not only bind growth factors and cytokines, but also retain water to promote ECM hydration and viscoelasticity^70^; thus, increased GAG retention may contribute to the increased contraction velocity and displacement observed in CMs cultured on CDMs. Sustained CM alignment and regimented contractile and electrophysiological training have been demonstrated to dramatically influence CM electrophysiology and contractility^71^, and it is therefore possible that extended culture of CMs on CDMs could improve their contractile function. However, changes in CM contractility also rely on CM plasticity as a factor of their differentiation and maturation stage, and further studies are required to determine the optimal CDM to support CM maturation.

In addition to structural differences, CDMs had protein compositions specific to their developmental stage. Fetal CDMs were enriched for the three primary chains of Col 6, the loss of which is associated with impaired skeletal muscle function and regeneration due to metabolic deficits ^62,63^. Col 6 is more abundant in the developing heart than in the mature heart^27^, but Col 6 deficiencies are not commonly associated with cardiac involvement except during periods of physiological stress, such as pregnancy^72^ or myocardial infarction^64^. The loss of Col 6 during infarction reduces myoactivation of resident CFs, thus ameliorating the pathological response and improving cardiac outcomes. However, such studies do not address whether cardiomyocytes, like skeletal myofibers^63^, rely on the presence of Col 6 for physiological respiration. Age-specific CDMs that recapitulated the fetal enrichment of Col 6 observed in cardiac tissue ECM provided the opportunity to directly assess the influence of Col 6 loss on CM function and metabolism. Fetal CDMs were more dependent on Col 6 for structural integrity than adult CDMs, and fetal CDMs that remained intact after Col 6 knockdown retained some residual Col 6 protein. In adult CDMs, loss of Col 6 had a detrimental impact on CM oxidative respiration without eliciting a compensatory increase in glycolysis, demonstrating the necessity of Col 6 for CM physiological respiration. Further exploration is required to conclusively determine the role of Col 6 in fetal hearts, but its abundance during development may serve to promote oxidative respiration in maturing cardiomyocytes, thereby facilitating the metabolic shift required for the increased ATP production^73,74^ in the oxygen-rich environment of the adult heart^75^. Notably, though, this metabolic deficit did not impair cardiac contractility or calcium handling, which is consistent with the lack of cardiac dysfunction in adult Col 6-deficient muscular dystrophies^65^. The differing metabolic and contractile responses further demonstrate that while the two functions are inherently linked in CMs by force-dependent bioenergetic demands^71,76^, they can be uncoupled by intermediary steps.

These studies demonstrate a significant advantage of CDMs, which is the ability to create, define and genetically alter *ex vivo* matrices that are compatible with multiple formats to meet diverse experimental requirements. The flexibility of the CDM model enabled longitudinal high-resolution microscopy during and after CDM formation and subsequent CM culture, as well as the ability to scale CDMs for paired transcriptomic/proteomic studies and the Seahorse XF96 flux assay. Further, this work underscores the utility of CDMs to study unique genetic and disease models without confounding cell type-specific differences in the secreted ECM (Fig. 1H). CDMs can reflect the composition and structural arrangement of ECM produced by various cell types without reflecting any potential homeostatic or paracrine influences of other tissues that would typically be impacted in a systemic disease state. Additionally, this approach can be adapted to produce CDMs from a variety of cell types, which will aid in distinguishing matrix-derived influences from autocrine/paracrine influences. CDMs that reflect the native cell-derived matrix composition will be useful for the study of ECM-related diseases involving multiple organs such as fibrosis, congenital disorders like Ehlers-Danlos and Marfan syndromes^77^, and aspects of developmental biology, where embryonic cells and their derivatives migrate through various ECM environments while simultaneously undergoing concurrent lineage specification^78^.

Overall, we demonstrate the development of a CDM model that enhances matrix production and retention without requiring the addition of exogenous matrix elements, thereby preserving many age- and tissue-specific ECM characteristics. Applying this method to cardiac biology, we identified complex interactions between CMs and cardiac ECM, including the contribution of Col 6 to CM mitochondrial respiration, but not contractility. These studies enable the dissection of complex cell-matrix interactions in tissue- and age-specific contexts to further clarify the impact of ECM on cell physiology and function throughout development.

## Materials and Methods

### Cardiac Fibroblast Culture

Primary human fetal and adult CFs were obtained from Cell Applications and maintained in Human Cardiac Fibroblast Media (Cell Applications, 316K-500). Mouse CFs were isolated from CD1 murine hearts at fetal (E14) or adult (8 week) ages via outgrowth selection. Briefly, heart were extracted from euthanized mice, minced, and allowed to settle onto tissue culture-treated plastic. Cells that migrated out of heart sections were collected as CFs. Mouse CFs were maintained in CF Media, comprised of DMEM (Corning, 15-013-CV) + 10% Fetal Bovine Serum (FBS; Atlanta Biologicals, S11150 lot M16139). Murine CFs were used to optimize dopa-mediated CDM retention (Fig. 1B) and MMC toxicity (Supp. Fig. 1). All CFs used for other applications were derived from human tissue.

### Cell-Derived Matrix Production

Prior to cell seeding, tissue culture multi-well plates were coated with 2mg/ml of poly L-dopamine hydrochloride (Sigma H8502) dissolved in a buffer of 100 mM Bicine (Sigma B3876), pH 8.5. Dopa stocks were stored under nitrogen at 4°C before coating. Plates were incubated in dopa solution overnight at room temperature (RT) with rotary agitation at 150rpm, and then rinsed vigorously with deionized water (diH_2_0). Plates were allowed to dry on the benchtop and were then sterilized by UV for 30 minutes, and stored at room temperature until use. CFs were seeded at confluency, between 50-60K/cm^2^ depending on well and cell size. Seeding was standardized to 55K (fetal) and 50K (adult) CFs in 24-well plates. Cells were cultured in CF media + 37.5mg/ml 70 kDa Ficoll (Fc; Sigma, F2878) and 25 mg/ml 400 kDa Ficoll PM (Sigma, F4375) (CF media + Fc) for 10 days with media completely refreshed every 2-3 days. Dextran (50kDa, MP Biomedicals, 0216011050) and polystyrene sulfonate (200kDa, Sigma, 561967) were introduced to CF media at 100μg/ml, respectively.

### Decellularization

Under non-sterile conditions, CDMs were decellularized by first rinsing with diH_2_0 twice, followed by 2 rounds of the following: 0.6M KCl for 1hr, 1M KI for 1hr, 2X diH_2_0 for 5 minutes at RT with rotary agitation at 150rpm. Matrices were then treated with 50u/ml DNase in 200mM Tris-HCl, pH 8.3 and 20mM MgCl_2_ for 1 hour at 37°C with rotary agitation, and then rinsed 2X with diH_2_0 for 5 minutes. CDMs were then either fixed for analysis, or transferred to a sterile environment and refreshed with PBS+/+ with 1X Penicillin-Streptomycin and fungizone at 0.25mg/ml and stored overnight at 4°C. The following day, CDMs were rinsed 2X with PBS and equilibrated at room temperature prior to seeding with iPSC-CMs.

### Mass Spectrometry

CDMs were rinsed overnight at 4°C in 0.5M NaCl in 10mM Tris at pH 7.5 + 1X cOmplete Protease Inhibitor Cocktail (PI), and subsequently solubilized with a 1% sodium dodecyl sulfate (SDS) solution in PBS + 1X PI at RT with agitation at 300rpm for 3-5 days until complete dissolution was achieved. Solubilized protein fractions were collected daily and replaced with fresh SDS + PI. Tris and SDS fractions were precipitated separately via chloroform/methanol separation and protein dry weight was quantified on an analytical scale. Dry protein was then solubilized in 8M Urea with 100mM Ambic and 10mM dithiothreitol (DTT), reduced with 500mM iodoacetamide in HPLC-grade water, and deglycosylated with 1000U PNGase F (P0704S, New England Biolabs). Solubilized protein was then digested with 3ug trypsin at 37°C overnight with agitation at 1400rpm before desalting on C18 columns. One microgram of the protein (determined by volume/original protein weight) was analyzed by LC-MS/MS. Unique peptides were identified by MaxQuant^79^ software referenced against the SwissProt human or mouse database. Data was subjected to Intensity Based Absolute Quantification (IBAQ)^80^ analysis and comparative analyses with statistical assessment using MSstats^81^. A total of 3 distinct CDMs/condition were analyzed. Pearson’s coefficient was calculated assuming a Gaussian distribution of 109 data pairs. The mass spectrometry proteomics data have been deposited to the ProteomeXchange Consortium via the PRIDE^82^ partner repository with the dataset identifier PXD020273. Identified proteins were cross-references to the Matrisome database (http://web.mit.edu/hyneslab/matrisome/)^45^ for the filtering of non-matrix proteins and for functional category assignments.

### Scanning Electron Microscopy

Decellularized CDMs were fixed with 2.5% (v/v) glutaraldehyde in 0.1 M sodium cacodylate and 0.03% aqueous eosin for 30 min with gentle agitation at room temperature. Fixed samples were washed with 0.1 M sodium cacodylate buffer, progressively dehydrated in an ethanol series and stored in absolute ethanol until critical point drying. Dried samples were immobilized on carbon tape, sputter-coated with gold and imaged by a specialized technician at Electron Microscopy Lab (UC Berkeley).

### Glycosaminoglycans

CDMs produced in 24 well plates with 0.1% dimethyl sulfoxide were rinsed with distilled water, incubated with agitation at 150 revolutions per minute in a 1% alcian blue solution in 3% acetic acid, pH 2.5, for 30 minutes at room temperature. Samples were rinsed once in dih_2_0 and imaged on a Keyence BZ-X710 microscope.

### Cardiomyocyte Differentiation

Human induced pluripotent stem cells (hiPSCs) (WTC11 cells modified with GCaMP6f reporter in the AAVS1 safe harbor locus; generously donated by Dr. Bruce Conklin) were seeded onto Matrigel-coated (80μg/mL; Corning, Corning, NY) plates at a concentration of 3×10^4^ cells/cm^2^ in mTeSR1 medium (Stem Cell Technologies, Vancouver, CA) supplemented with 10μM ROCK inhibitor (Y-27632, SelleckChem, Houston, TX) for 24h. Differentiation of hiPSCs to CMs was performed using a serum-free, chemically defined protocol. Briefly, once hiPSCs reached 100% confluence (~3-4 days; denoted as differentiation day 0), cells were fed with 12μM CHIR (SelleckChem) in RPMI1640 medium (Thermo Fisher, Waltham, MA) with B27 supplement without insulin (RPMI/B27-; Life Technologies, Grand Island, NY). After 24h, CHIR was removed by feeding with RPMI/B27- and on day 3, cells received a 48h-treatment with 5μM IWP2 (Tocris, Bristol, UK) in RPMI/B27-. After 7 days, the culture medium was switched to RPMI1640 containing B27 supplement with insulin (RPMI/B27+; Life Technologies) and completely refreshed every 3 days thereafter. After 15 days of differentiation, hiPSC-CMs were dissociated with trypsin and re-plated at 50K/cm^2^ onto CDMs in RPMI/B27+ supplemented with 10μM ROCK inhibitor and 10% FBS. Cultures were refreshed with RPMI/B27+ media after 2 days, and cultured for 1-4 days prior to functional assessment and/or fixation with 4% paraformaldehyde (PFA) for 15 minutes and 3X rinses with Phosphate Buffered Solution (PBS).

### Immunofluorescent Microscopy

Proteins were visualized by immunocytochemistry, wherein fixed CDMs and/or cells were blocked in 5% normal donkey serum (017-000-121, Jackson Immunoresearch Labs) and 0.3% Triton-X 100 (T8787, Sigma) in PBS for 1 hour at RT. Antibody dilution buffer (Ab Buffer) was comprised of PSB supplemented with 1% BSA (A4503, Sigma) and 0.3% Triton-X 100. Samples were incubated with primary antibodies in Ab Buffer for 1 hour at RT or overnight at 4°C, followed by 3 washes with PBS and 1 hour of incubation with secondary antibodies at 1:300 in Ab Buffer at RT and 3 subsequent PBS washes. Primary antibodies and stains were used as follows: cardiac troponin T (Abcam ab45932, 1:400), fibronectin (Sigma F6140, 1:400), Plakophilin 2 (VWR 101981-232, 1:10), Col 6 (Abcam ab6588, 1:500), Hoescht 33342 (ThermoFisher 62249, 1:10,000), Phalloidin (ThermoFisher A34055, 1:100). Samples were imaged on a Zeiss Z1 Inverted or Zeiss LSM880 Confocal Microscope.

### CM Contraction Dynamics

CM contraction and calcium handling properties were quantified from recorded videos of CMs maintained at in Tyrode’s solution (137□mM NaCl, 2.7□mM KCl, 1□mM MgCl_2_, 0.2□mM Na_2_HPO_4_, 12□mM NaHCO_3_, 5.5□mM D-glucose, 1.8□mM CaCl_2_; Sigma–Aldrich) at physiological conditions (37°C, 5% CO_2_, OkoLab). Cells were placed into Tyrode’s solution 1 hour prior to data acquisition and maintained in a 37°C incubator until imaging. CM contraction frequency and speed were assessed from recorded videos captured with brightfield illumination and assessed with the Pulse contractility analysis (Dana Solutions). Optical flow gradients of CM contractions were quantified by custom python scripts. Calcium handling properties were quantified by recording from fluorescence intensity of the genetically encoded GCaMP6f calcium indicator. Stroke velocities were calculated using custom python scripts. Source code is available at: https://github.com/david-a-joy/multilineage-organoid^83^. A total of 4 experimental replicates were measured per condition, and video recordings were performed at 50ms (brightfield) or 10ms (fluorescence) exposure times through Zen software on a Zeiss Z1 Inverted Microscope.

### Western Blot

CFs subjected to transfection with NT or anti-Col 6 siRNA were incubated for 10 min on ice in RIPA Buffer (Sigma-Aldrich). Samples were then centrifuged at 13,000xg for 15 minutes and supernatants were collected. Protein concentrations were determined using a Pierce BCA Protein Assay kit (Thermofisher Scientific, 23225) colorimetric reaction and quantified on a SpectraMax i3 Multi-Mode Platform (Molecular Devices). Protein concentrations were determined by Bradford assay (Thermo Fisher, 23225). Protein from each lysate (17.5μg) was suspended in 1X Sample buffer (BioRad) containing beta-mercaptoethanol, boiled for 5 minutes and separated on precast 12% SDS-PAGE TGX gels (BioRad, 465-1044). Protein gels were transferred to a nitrocellulose membrane, blocked in 1:1 PBS: Blocking Buffer (Rockland Inc, MB-070) for 1hr at room temperature, and subsequently incubated at 4°C overnight with primary antibodies anti-Col 6 (Abcam, ab182744, 1:2000) and anti-GAPDH, (Invitrogen 1:1,000). Membranes were washed with 0.1% Tween in PBS, followed by incubation (45 min at room temperature) with infrared secondary antibodies: IRDye 800CW and IRDye 680CW (LI-COR, 1:5,000) and imaged on an ChemiDoc MP Imaging System (BioRad). Band intensity was analyzed in Image Lab v2.4 (BioRad). Representative CDMs for each condition were assessed by western blot.

### Col 6 Knockdown

Col 6 KD in CDMs was achieved by serial lipid-mediated transient transfections of pooled siRNAs against COL6A1, Col6A2 and Col6A3 (Dharmacon, M-011620-01, M-011621-01, M-003646-02) or a non-targeting control (Dharmacon, D-001210-01) in CFs during CDM production every 3-4 days. Complete CF media was refreshed at least 1 hour prior to transfection. siRNAs were diluted to a final concentration of 100nM in Opti-MEM Reduced Serum Medium (ThermoFisher, 31985070) before mixing with an equal volume of Optimem:RNAiMax (ThermoFisher, 13778150) at a ratio of 50:1. The siRNA:RNAiMax mixture was complexed at room temperature for 5 minutes before adding to cultures dropwise. Media was refreshed 6-18 hours later with CF media + Fc, and then cultured normally. In 24 well plates, media was refreshed with 400μl CF media and transfected with a final volume of 100μl siRNA:RNAiMax.

### Seahorse Assay

For metabolic assays, fetal and adult CFs were seeded onto dopa-coated Seahorse XF96-well cell culture microplates (V3 PS, Agilent, 101085-004) at a density of 123k/cm^2^. XF96 plates were refreshed with 60μl CF media for 1 hour prior to transfections, and then 20μl siRNA:RNAiMax pre-diluted with 20ul DMEM (Corning, 15-013-CV) was introduced to each well drop-wise, following a 5 minute pre-incubation step. Transfections were repeated every 3-4 days prior to decellularization at day 10. The following day, iPSC-derived cardiomyocytes between days 12 and 15 of differentiation were dissociated with Trypsin supplemented with 10μM ROCK-Inhibitor for 30 min and seeded onto the decellularized Seahorse plate in RPMI/B27+ supplemented with 10μM ROCK inhibitor and 10% FBS at a density of 439k/cm^2^ and cultured for 3 days. Before the metabolic assay, media was refreshed with Seahorse XF RPMI medium, pH 7.4 (Agilent, 103576-100) supplemented with 10 mM Glucose (Agilent, 103577), 2 mM Sodium Pyruvate (Agilent, 103578), and 1 mM Glutamine (ThermoFisher, 35050061) pH 7.4, and incubated for one hour at 37°C without CO_2_. Oxygen consumption rate (OCR) and extracellular acidification rate (ECAR) were measured using the Seahorse XF96 Extracellular Flux Analyzer (Agilent) during repeated cycles of 3x 3 minutes mix, 3 minutes measure, after sequential injection of the following mitochondrial inhibitors: ATP synthase inhibitor oligomycin (final concentration: 1.5 uM), proton ionophore fluorocarbonyl cyanide phenylhydrazone (FCCP: 1 uM), and complex I inhibitor rotenone/Antimycin A (0.5 uM). Metabolic parameters were analyzed using Wave software version 2.6.1 and normalized per 10^3^ iPSC-CMs. A total of 2-8 CDM replicates were analyzed per CDM condition, depending on CDM integrity, which was assessed prior to CM seeding.

### RNA Sequencing

Cultured CFs were lysed with Trizol (Ambion), and all groups were collected in triplicate. RNA was extracted using the RNeasy Mini Kit (Qiagen 74104) and quantified using the NanoDrop 2000c (ThermoFisher Scientific). RNA-seq libraries were created using the SMARTer Stranded Total RNA Sample Prep Kit (Takara Bio) and sequenced on the NextSeq 500 (Illumina) to a minimum depth of 25 million reads per sample. Initial reads were aligned to GRCh37 human genome using HiSAT2 2.1.0^84^. Genes with low counts were filtered and gene expression levels normalized across samples using edgeR 3.28.1^85^. Counts were transformed to log2-counts per million (logCPM) using limma-voom 3.42.2^86^. Significantly different genes were assessed at a Benjamini & Hochberg adjusted p < 0.05 level and a minimum logCPM absolute difference of 1 between the gene and the mean expression level. GO term analysis was performed on significantly up and downregulated genes using enrichGO from clusterprofiler 3.14.3. Raw data will be available at GEO (Accession #:GSE155225).

### Statistical analysis

All results are shown as mean ± standard deviation. Unless otherwise stated, all statistical analysis of the data was performed using Prism 8 with statistical significance determined at p < 0.05 with a one way ANOVA and subsequent Dunnett’s test to correct for multiple comparisons.

## Acknowledgements

This work was supported by the California Institute of Regenerative Medicine Grant LA1-08015 (T.C.M) and NIH P01HL089707 (N.J.K). M.A.K. was supported by an American Heart Association Predoctoral Award (17PRE33660113) and a QB3 Frontiers in Medial Research Fellowship. D.A.J. was supported by a UCSF Genentech Fellowship. The authors would like to thank the Stem Cell Core (Dr. Po-Lin So; Roddenberry Stem Cell Foundation), Histology and Light Microscopy Core (Dr. Meredith Calvert), and Genomics Core (Dr. Natasha Carli and Jim McGuire) at the Gladstone Institutes, and Electron Microscopy Lab (Dr. Danielle Jorgens) at UC Berkeley for all the help and expertise shared, as well as Dr. Bruce Conklin’s laboratory for providing WTC11 GCaMP iPS cell line. The authors are also grateful to Phil Messersmith and Devang Amin, of UC Berkeley, for their guidance and support in adapting L-polydopamine for cell culture applications. A special thanks to Drs. Tracy Hookway and Marsha Rolle for relevant discussions, and to Oriane Matthys for technical assistance and discussion. The authors also acknowledge the Gladstone Scientific Editing Department (Kathryn Claiborn) for reviewing the manuscript and scientific editing.

## Author Contributions

M.A.K. and T.C.M. conceived of and designed the experiments; M.A.K., S.J.R., N.M.C., A.C.S. and M.N.W. performed the experiments; D.A.J. developed the python script for calcium imaging analysis and performed alignment, normalization and statistical evaluation of RNASeq data; E.S. performed mass spectrometry experiments and D.L.S. performed proteomic IBAQ and MSstats analyses; D.L.S. and N.J.K provided oversight for proteomic experiments; A.C.S. designed the figure schematics; M.A.K. and T.C.M. wrote and edited the manuscript.

## Supplemental Figures

**Supplemental Figure 1.**
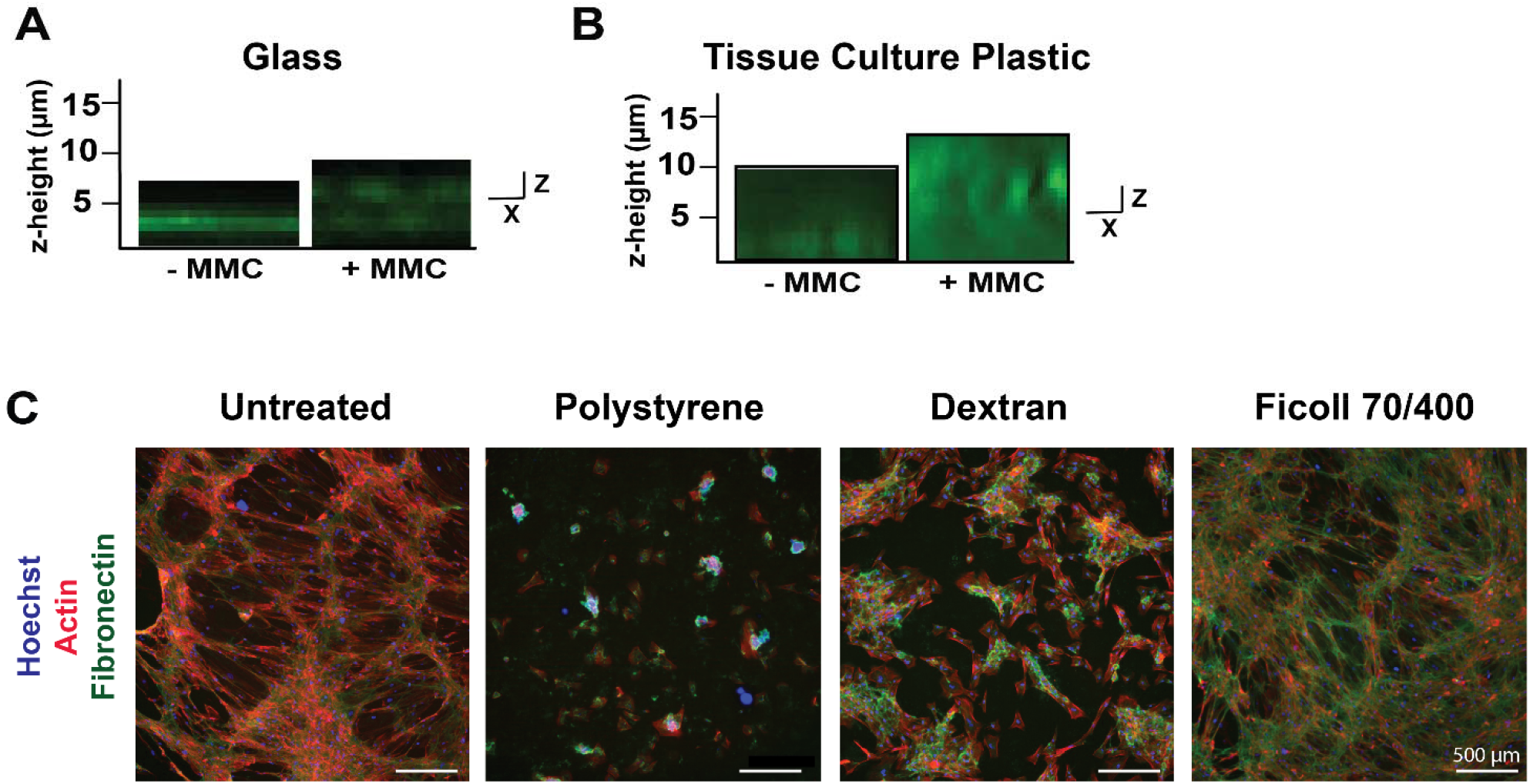
A-B) Optical sectioning of human CDMs produced with and without MMCs on glass (A) and tissue culture plastic (B). C) Mouse CFs cultured with various MMCs.

**Supplemental Figure 2.**
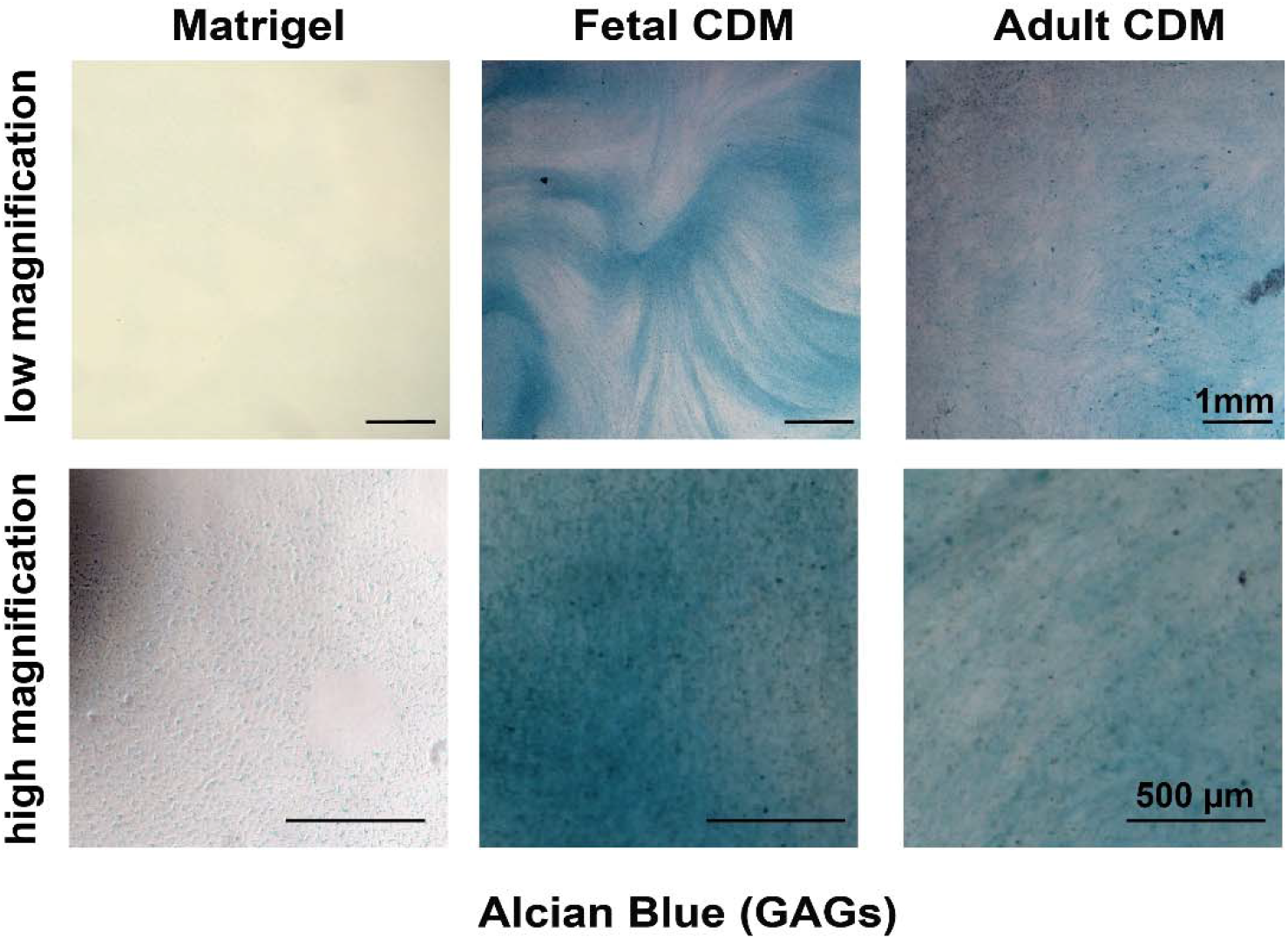
GAGs present in Matrigel and human CDMs, detected by alcian blue staining. Low magnification (top) and high magnification view (bottom).

**Supplemental Figure 3.**
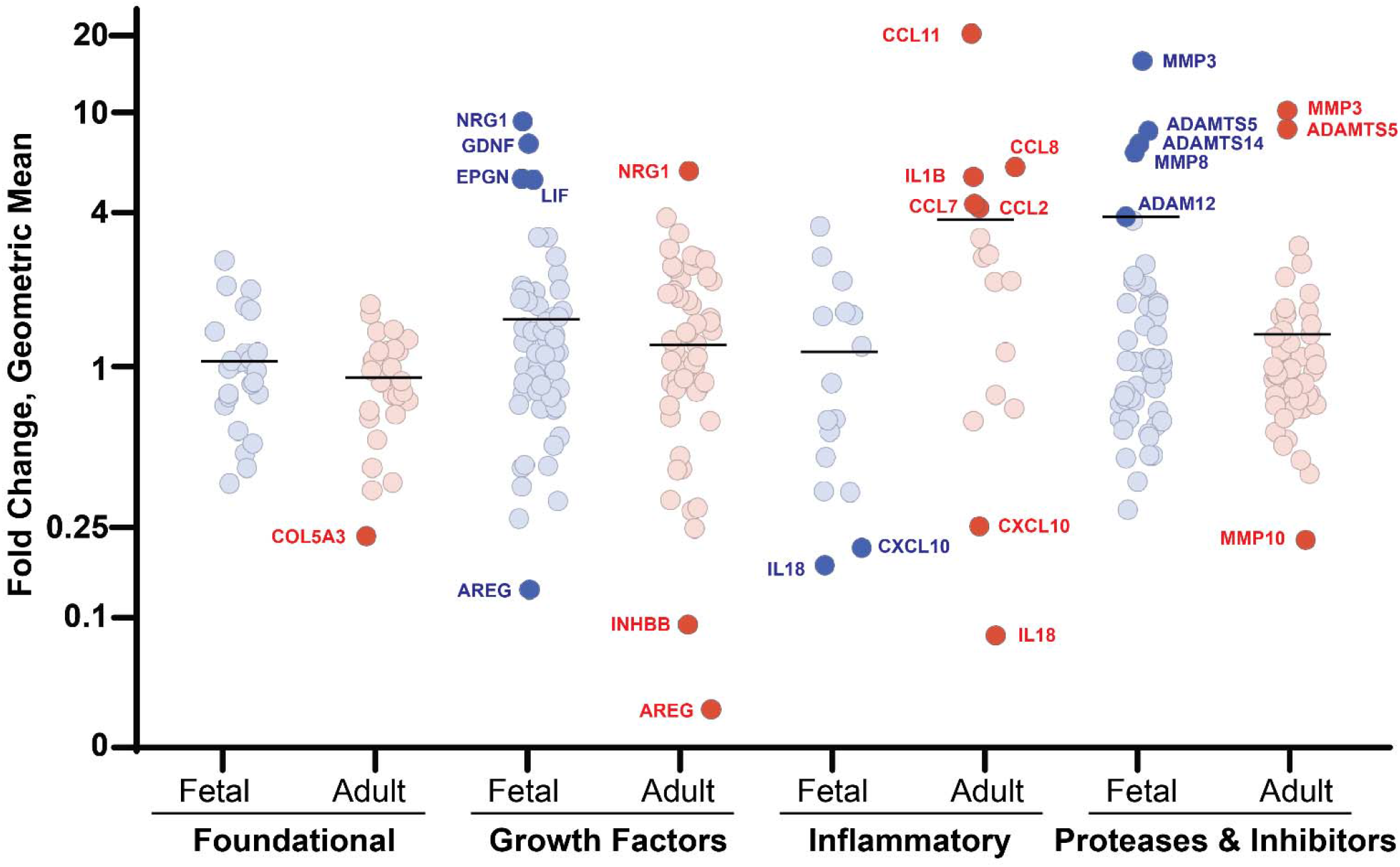
Geometric mean of matrix transcript fold changes assessed at days 1, 5 and 10 of human CF culture, designated by functional category. Highlighted genes have transcripts more than 4-fold up or down regulated from day 1 expression.

**Supplemental figure 4.**
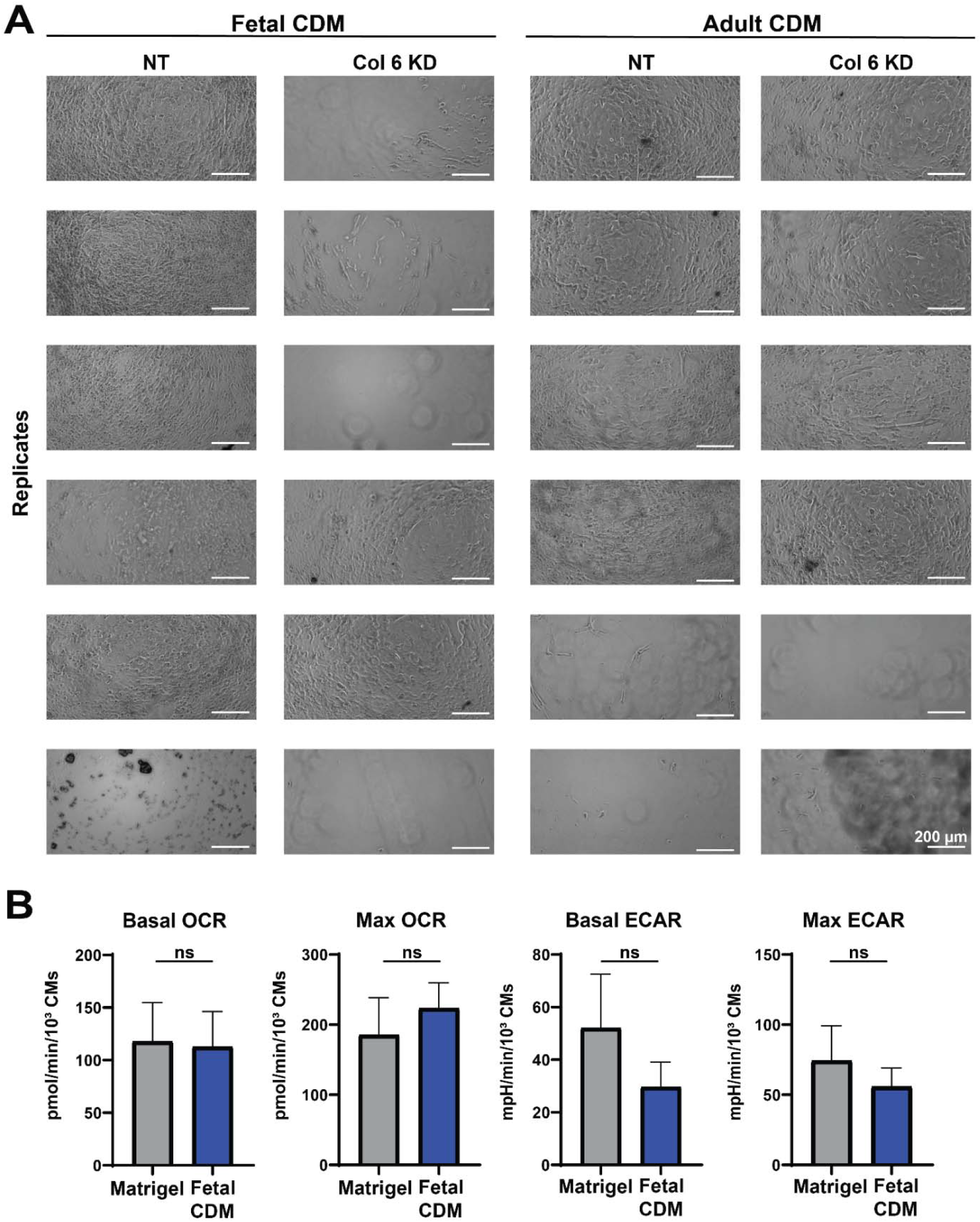
A) Brightfield images of human CDMs in Seahorse XF 96 plate. CDM destabilization with Col 6 KD was increased in fetal CDMs (from 1 NT CDM to 4 KD CDMs) but not adult CDMs (from 2 NT CDMs to 2 KD CDMs). B) Quantification of basal (baseline) and maximal (FCCP-treated) OCR and ECAR of iPSC-CMs cultured on Matrigel and fetal CDMs show no significant differences. ns = not significant.

